# Changes in salt-marsh vegetation weakly affect top consumers of aquatic food webs

**DOI:** 10.1101/2020.07.27.222406

**Authors:** Lafage Denis, Carpentier Alexandre, Sylvain Duhamel, Christine Dupuy, Eric Feunteun, Sandric Lesourd, Pétillon Julien

## Abstract

Salt marshes are under high, and increasing, anthropogenic pressures that have notably been reported to affect the diet of several fish species, probably resulting in nursery function alterations. Most of the previous studies in Europe were yet based on gut content analysis of fish, which can be considered a snapshot of immediate impacts of salt-marsh changes, and hardly of long-term effects of disturbances. In this study, we investigated the impact of vegetation type (resulting from both plant invasion and sheep grazing) by assessing trophic network (and especially fish diet and position) of different salt-marsh conditions. Replicated samples of basic sources (particular organic matter and microphytobenthos), dominant vegetation, potential aquatic and terrestrial prey and fish of 3 main species were taken during summer 2010 in two bays from Western France (Mont -Saint-Michel Bay and Seine Estuary) and analysed using C and N stable isotope compositions. All response variables tested (overall trophic organization, trophic niche and trophic position) provided consistent results, i.e. a dominant site effect and a weaker effect of vegetation type. Site effect was attributed to differences in anthropogenic Nitrogen inputs and tidal regime between the two bays, with more marine signatures associated with a higher frequency of flooding events. A second hypothesis is that *E. acuta*, which has recently totally replaced typical salt-marsh vegetation in Mont Saint-Michel Bay strongly impacted the nursery function. The trophic status of dominant fish species was unchanged by local salt-marsh vegetation, and considered consistent with their diet, i.e. high for predatory species (the sea bass *Dicentrarchus labrax* and the common goby *Pomatoschistus microps*) and lower for biofilm grazing species (the thinlip mullet *Chelon ramada*). This study finally highlights the relevance of stable isotopes analyses for assessing long-term and integrative effects of changes in vegetation resulting from human disturbances in salt marshes.

**Highlights:**

- Cross-ecosystem subsidies are of high functional importance, notably in salt marshes
- Fish are vectors of exchanges, most European studies being based on their gut content
- Using stable isotopes we analysed the effect of surrounding vegetation on food webs
- Surprisingly we found weak vegetation and strong site effects on all metrics
- Nitrogen inputs, site accessibility and loss of nursery function can explain this fact

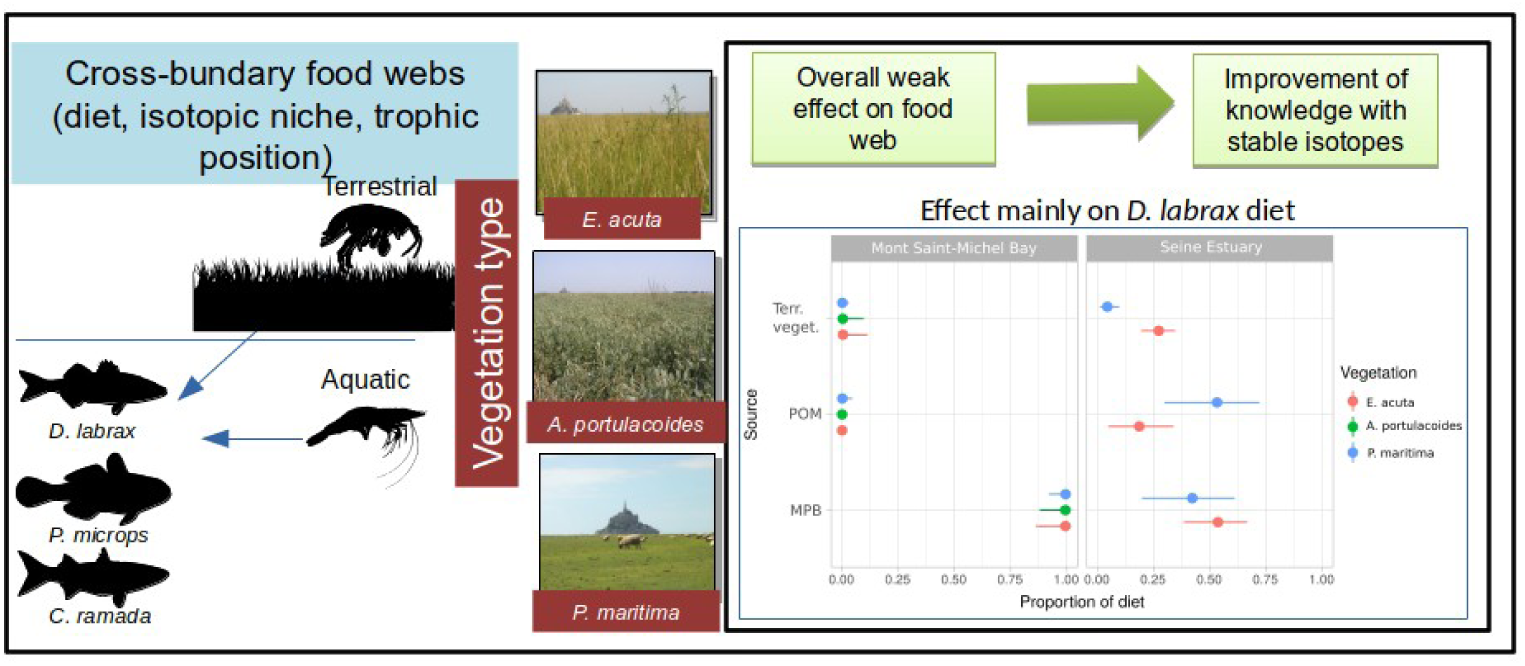

## 1. Introduction

Mostly localized in the vicinity or within estuaries, salt marshes have typical ecotone functioning with influences from continental freshwaters and coastal marine waters together with anthropic pressures from both origins (McLusky and Elliott, 2004). This particular location explains, for instance, the very high primary production (up to 20t of dry matter per ha and per year: Lefeuvre et al., 2000), additional nutrients being brought to these habitats, coming either from catchments or from the sea. Conversely, organic matter resulting from the decomposition of plant material is partly exported to the sea after high tides, as an outwelling process according to Odum (1980). Located in intertidal areas, salt-marsh vegetation is covered in Europe by tides only one or two times a month in macrotidal areas. Nevertheless, marine waters drill webs of drainage creeks permitting temporary access to aquatic fauna to salt marshes resources during the flood. The high primary production leads to important populations of arthropods and notably decomposers, living within and eating on litter of halophilic plants (Schrama et al., 2012). Some of these arthropods are in turn largely consumed by insectivorous birds (Geslin et al., 2006) and by juveniles of fish when they fall into creeks during flooding (Joyeux et al., 2017; Laffaille et al., 1998). Consumers, when feeding upon resources from salt marshes, are thus likely to contribute, indirectly, to the export of organic matter from salt marshes to adjacent sea waters (Laffaille et al., 2002; Lefeuvre et al., 1999). The complex created by salt marshes and their creeks thus plays a key role for the survival of many fish species during their first year, and can be qualified as a nursery (e.g. Cattrijsse et al., 1994; Kneib, 1997, the review of Cattrijsse and Hampel, 2006; Green et al., 2012, 2009; Joyeux et al., 2017; Nunn et al., 2016). This particular functioning can be modified by many disturbances, especially when salt-marsh vegetation is submitted to numerous pressures and among them, plant invasion and sheep grazing.

Biological invasions can indeed have huge consequences on ecosystems (see Grosholz, 2002 for review), being able to affect biogeochemical pool and fluxes of materials, and thus structure and functioning of the entire trophic web (Charles and Dukes, 2007; Ehrenfeld, 2010; Pejchar and Mooney, 2009 for review). Indeed, plants play a major role in habitat functioning notably through their physical structure, their cover and/or particular characteristics, which often determine the physical characteristics of habitats (Ellison et al., 2005). In this context, invasions by plants are reported as a disturbing factor in several salt marshes but remain obviously less studied compared to terrestrial and freshwater habitats (Grosholz, 2002). Most of these studies dealt with *Spartina spp*. (Ayres et al., 2004), with changes in habitat structure and cascading effects on birds (Gan et al., 2010; Ma et al., 2011), arthropods (Wu et al., 2009), benthic macrofauna (Hedge and Kriwoken, 2000; Neira et al., 2006), diet of deposit-feeding snails (Wang et al., 2014) and global carbon and nitrogen fluxes and stocking (Chen et al., 2011; Cheng et al., 2008; Davidson et al., 2018; Yang et al., 2016; Yin et al., 2015). Other examples include *Tamarix spp*. invasion and its effect on marsh consumers (Whitcraft et al., 2008) or invasion by *Phragmites australis* driving the loss of plant diversity (Silliman and Bertness, 2004). An atypical case of invasion was also reported for the nitrophilous grass *Elytrigia acuta* (formerly *Elymus athericus*), particularly spectacular in the Mont-Saint-Michel Bay (Valéry et al., 2017).

In the absence of invasive species, plant composition of salt marshes is first driven by inundation frequency and resulting gradient in soil salinity (e.g. Pétillon et al., 2010), but also depends largely on management practices (Bouchard et al., 2003). Inundation frequency and management practices will both affect on-going processes and consequently higher trophic levels. As an example, intensively grazed or mown salt marshes tend to be favourable to herbivore birds such as the brant goose *Branta bernicla* or the graylag goose *Anser anser* (*Val*éry et al., 2008), because of dominance by short stands of *Puccinellia maritima* grass. On the opposite, the absence of management will increase habitats with *Atriplex portulacoides* or the invasive *Elytrigia acuta*, with higher arthropod densities (Pétillon et al., 2005), and more insectivorous birds (Geslin et al., 2006). The impact of salt-marsh management is, for instance, obvious on potential prey of fish (e.g. Ford et al., 2012;

Pétillon et al., 2007; van Klink et al., 2014). Salt marshes are indeed essential habitats for particular stages of life cycles from numerous species like fish or invertebrates (Boesch and Turner, 1984; Kneib, 1997; Laffaille et al., 2001; Rountree and Able, 1992). Nursery function seems particularly impacted by habitat disturbance, and notably by changes in dominant vegetation (Dionne et al., 1999; Joyeux et al., 2017; Laffaille et al., 2005, 2000; Warren et al., 2002). Sheep grazing, especially at a high density of livestock, strongly affects local biodiversity (Leroy et al., 2014), with probably stronger deleterious effects than the invasive plant (Puzin and Pétillon, 2019), and potentially affects fish nursery as revealed by gut content analyses of few fish species (Laffaille et al., 2001).

Plant community is often dominated by a single or few plant species in salt marshes which can thus be considered as keystone species affecting other species and/or matter fluxes out of proportion to their abundance and through entirely biotic mechanisms (Power et al., 1996). In this study, we tested the broad hypothesis that changes in salt-marsh vegetation composition (here resulting from plant invasion and grazing) cascade into aquatic food webs, including their top-consumers i.e. dominant fish species (see e.g. Laffaille et al., 2005; Mantzouki et al., 2012). More precisely, the following expectations were tested: (i) a strong impact of vegetation and a lesser site effect on both fish diet and trophic niche, and (ii) some changes in trophic niche and position only visible for predatory species. These hypotheses were tested by comparing the C and N stable isotope compositions of trophic networks from invasive, grazed and natural vegetation from two bays of Western France. These sites being spatially quite close each to other, we had no particular expectation of a strong site effect.

## 2. Methods

### 2.1. Site description

Two salt marshes located north-west of France were sampled: the Mont Saint-Michel Bay and the Seine Estuary (Fig .1). The Mont Saint-Michel Bay is located at the border between Brittany and Normandy at North-western France (latitude 48°40’N, longitude 1°31’W). It is a macrotidal bay characterized by one of the highest tidal range in the world (15m). Intertidal area covers 220 km^2^, composed of 180 km^2^ of mudflats (slikke) and 40 km^2^ of saltmarshes (schorre). Saltmarshes are drained by a web of creeks filled by tidal waters when the tidal range exceeds 11.25m (Lefeuvre et al., 2000) and during 2 to 3 hours according to the water level. Contrary to the Mont-Saint-Michel Bay, the Seine Estuary has been heavily transformed by human notably to reduce tidal influence on intertidal habitats. It is located in the middle of the coast in the Normandy region (latitude 49°27’ N, longitude 0°16’E). The tidal range reaches 7m. Intertidal area covers 16 km^2^ and is composed of 4 km^2^ of mudflats (slikke) and 12 km^2^ of saltmarshes (schorre). Sampling sites were located in the middle estuary, which is characterized by a strong salinity gradient (Fig. 1). Like in the Mont-Saint-Michel Bay, tidal waters filled the creeks during the flow when tidal range exceeds 3 m and during 5h according to the water level.

**Figure 1:**
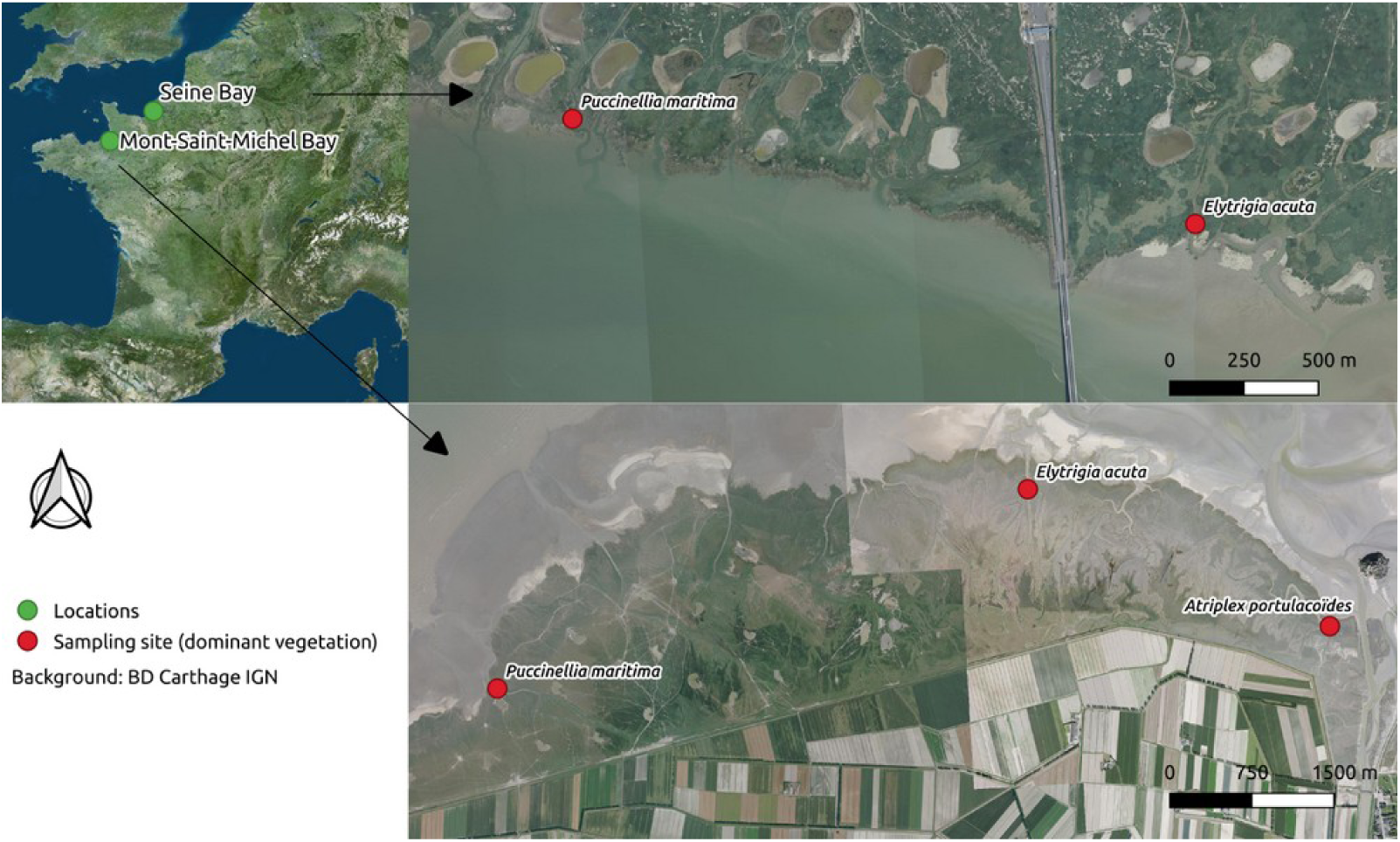
Sampling site location.

Two sites were sampled in the Seine Estuary: one was dominated by grazed *Puccinella maritima*, the other by invasive *Elytrigia acuta*. In the Mont-Saint-Michel Bay, three sites were sampled: one was dominated by ungrazed *Puccinella maritima*, the second by non-invasive stands of *Elytrigia acuta* and the last one by *Atriplex portulacoides*.

### 2.2. Food web sampling and stable isotopes preparation

Sampling took place during summer 2010. A fykenet (4 mm mesh size) was set across one representative creek within each site in both salt marshes to catch fish leaving sites after the ebb (see Joyeux et al., 2017) for more details on a similar design). All fish captured were identified and measured to get a proxy of individual age. Only most abundant species, i.e. sea bass (*Dicentrarchus labrax*), thinlip mullet (*Chelon ramada*) and common goby (*Pomatoschistus microps*) were kept for isotope analyses. Size ranges were 30.83 ± 8.40 mm and 26.50 ± 5.30 mm for sea bass, 19.96 ± 4.82 mm and 20.10 ± 4.90 for common goby, 55.60 ± 11.1mm and 72.5 ± 12.6 mm for thinlip mullet in Mont Saint-Michel Bay and Seine estuary, respectively. White muscle tissues were collected from each individual at the laboratory.

Potential prey were collected by hand or caught in the fykenet when sampling fish. A minimum of 3 samples per site and per predator or potential prey species (based on previous studies using gut content identification, e.g. (Laffaille et al., 2005, 2002, 2001) was collected. For species submitted to decarbonation, a fourth sample (not submitted to decarbonation) was collected to check for the possible impact of the decarbonation treatment on δ^15^N values (Pinnegar and Polunin, 1999). Particulate organic matter (POM), microphytobenthos (MPB) and terrestrial vegetation were sampled as different basic sources. Terrestrial vegetation was collected by hand. POM came from 1L of water collected in the creak during the ebb. The water was filtered on decarbonated (4h at 550°C) GF-F filters (47 mm diameter Wathmann).

MPB was extracted from the superficial layer (5mm) of scrapped muddy sediment by exposing it to light during 2 hours. The sediment was then covered with a 100 μm nylon filter and sand previously sieved (63 et 250 μm) and carbonated (5 hours, 550°C). After waiting several hours for MPB to migrate to the surface through the nylon filter, the superficial layer (2mm) was scraped and sieved under seawater on a 45 μm filter. The content of the filter was then filtered again under decarbonated (4h, 550°C) GF-F filters.

Except for filters (POM and MPB), which were dried during 24h at 50°C in a desiccator, all samples were then frozen before being lyophilized and crushed into a fine powder.

No delipidation was performed on lyophilized samples since no significant effect on isotope values was found during preliminary tests (cyclohexane treatment according to DeNiro and Epstein, 1977). This choice was also based on the C/N threshold of 3.5 proposed by Post et al. (2007) for aquatic animal samples.

### 2.3. Stable isotopes analyses

The nitrogen and carbon isotopic compositions were determined using EA-IRMS (Isoprime, Micromass, UK). The carbon and nitrogen isotope ratios are expressed in the delta notation δ^13^C and δ^15^N where: δX = [(R_Sample_ / R_Reference_) − 1] × 1000, where X = δ^13^C or δ^15^N and R is the ratio ^13^C: δ^12^C or ^15^N: ^14^N in the sample and in the reference material. Results are referred to Vienna Pee Dee Belemnite (VPDB) for C and to atmospheric nitrogen for N and expressed in units of ‰ ± standard deviation (sd). The analytical precision (standard deviation for repeated measurements of internal standards) was ± 0.2 ‰ and ± 0.3 ‰ for δ^13^C or δ^15^N, respectively.

### 2.4. Statistical analyses

Isotopic signatures of fish, preys and baselines were compared using MANOVA analysis with site, vegetation and site/vegetation interaction as factors.

Fish diet partitioning was achieved using Bayesian mixing model performed with package MixSIAR 3.1.10 (Stock and Semmens, 2013) in R 3.6.1 (R Core Team, 2020) with two tracers: δ^13^C and δ^15^N.

As a first step, models were built for each fish species with POM, MPB and terrestrial vegetation as potential sources. We used baseline sources rather than potential preys as we were interested in studying the effect of terrestrial vegetation changes in the overall contribution of salt-marsh components. For species with a significant effect of vegetation changes, we built models with potential preys (inferred from stomach contents).

For each fish species, one null model (no random or fixed effect) and models including different combinations of random (site) and fixed (vegetation) effects were tested with the following settings: 3 chains with 300 000 iterations and burn-in of 200 000 to allow for adequate model convergence. Site was included as a random factor for both predator and source signatures. Model convergence was checked using Gelman-Rubin and Gewek diagnostics. Candidate models were then compared using leave-one-out cross-validation (LOO). Input values for each model were the raw isotopic values for fish and mean and standard deviation of each source. To compare the relative contribution of factors included in the best model, a new model with both factors included as random variables was run. Trophic enrichment factor values were derived from Kostecki et al. (2012) following Selleslagh et al. (2015). Two sets of values were used according to fish species trophic guilds. Values for predators (*D. labrax* and *P. microps*) were: 2 ± 0.6% and 5.6 ± 1.5% for δ^13^C and δ^15^N respectively. Values for grazers (*C. ramada*) were: 1 ± 0.6% and 3.4 ± 1.5% for δ^13^C and δ^15^N respectively (Carpentier et al., 2014; Como et al., 2018).

The fish trophic position was estimated using a two sources Bayesian model. The two sources included were the ones contributing most to the fish diet according to previous mixing models. Estimation was performed using the tRophicPosition 0.7.7 package (Quezada-Romegialli et al., 2019). Input values for each model were the raw isotopic values for fish and sources. The grouping variable was vegetation type.

The isotopic niche of fish was approximated through standard ellipse area derived from δ^13^C and δ^15^N values using the Bayesian framework developed by Jackson et al. (2011) and implemented in the R package SIBER 2.1.4. The standard ellipse area estimation requires at least 5 observations per modality (here per site and vegetation type), which prevented us from including *D. labrax* and *P. microps* sampled in *A. portulacoides* in the Mont-Saint-Michel Bay.

## 3. Results

### 3.1. Overall stable isotope composition of salt-marsh food web

The MANOVA on fish isotopic signatures (Table 1, Appendix A) demonstrated a significant effect of fish species (F=49.23, df=4, p<0.001), site (F=220.86, df=2, p<0.001) and vegetation type (F=9.07, df=4, p<0.001), but no interaction between site and vegetation type (F=2.48, df=2, p=0.087).

**Table 1:** Mean and standard deviation of isotopic values

|  |  | Mont-Saint-Michel Bay |  | Seine Estuary |  |
| --- | --- | --- | --- | --- | --- |
| | Species | Mean $\delta^{13}\text{C} \pm \text{sd}$ | Mean $\delta^{15}\text{N} \pm \text{sd}$ | Mean $\delta^{13}\text{C} \pm \text{sd}$ | Mean $\delta^{15}\text{N} \pm \text{sd}$ |
| Fish | *Dicentrarchus labrax* | -15.69 ± 0.48 | 13.43 ± 0.42 | -18.81 ± 1.06 | 14.87 ± 0.61 |
| | *Pomatoschistus microps* | -15.32 ± 0.67 | 13.77 ± 0.6 | -18.86 ± 1.74 | 16.31 ± 0.6 |
| | *Chelon ramada* | -16.53 ± 1.9 | 10.42 ± 0.71 | -19.79 ± 3.24 | 13.85 ± 1 |
| Aquatic prey | *Carcinus maenas* | -16.2 ± 1.22 | 10.07 ± 0.68 | -17.52 ± 1.56 | 12.11 ± 0.64 |
| | *Hediste diversicolor* | -16.12 ± 0.67 | 8.98 ± 0.84 | -17.14 ± 0.08 | 12.09 ± 0.49 |
| | *Lekanesphaera rugicauda* | -18.12 ± 2.22 | 7.39 ± 0.78 | -19.36 ± 2.34 | 8.28 ± 0.77 |
| | Mysida | -15.81 ± 0.09 | 11.04 ± 0.18 | -18.41 ± 0.56 | 12.17 ± 0.26 |
| | *Palaemonetes varians* | -18.38 ± 1.13 | 10.86 ± 0.82 | -19.51 ± 0.42 | 13.49 ± 0.99 |
| | *Corophium volutator* | | | -20.55 ± 0.61 | 9.23 ± 0.77 |
| Terrestrial prey | *Chortipus albomarginatus* | -24.73 ± 0.57 | 7.18 ± 0.95 | -23.54 ± 0.23 | 9.42 ± 0.62 |
| | Homoptera | -23.56 ± 0.17 | 6.09 ± 0.1 | -24.38 | 6.44 |
| | Lycosidae | -23.37 ± 0.64 | 12.68 ± 1.31 | -24.05 ± 0.18 | 14.31 ± 0.6 |
| | *Orchestia gammarellus* | -23.43 ± 0.89 | 7.37 ± 1.64 | -23.23 ± 0.7 | 10.67 ± 0.37 |
| | *Pogonus chalceus* | -24.39 ± 1.27 | 11.35 ± 0 | -23.9 ± 1.4 | 17.8 ± 1.39 |
| Terrestrial |  |  |  |  |  |
| vegetation | *Atriplex portulacoides* | -25.42 ± 1.77 | 8.88 ± 0.63 | | |
| | *Puccinellia maritima* | -24.83 ± 1.21 | 6.06 ± 0.8 | -25.84 ± 1.9 | 8.56 ± 0.72 |
| | *Elytrigia acuta* | -24.73 ± 1.11 | 6.22 ± 1.09 | -24.73 ± 2.76 | 6.73 ± 1.05 |
| | *Salicornia spp.* | -26.3 ± 0.71 | 7.02 ± 0.29 | -27.21 ± 1.07 | 8.17 ± 1.4 |
| MPB | | -17.07 ± 1.2 | 6.9 ± 0.41 | -19.75 ± 0.66 | 10.92 ± 1.04 |
| POM | | -20.37 ± 1.17 | 4.09 ± 1.09 | -24.42 ± 1.15 | 9.4 ± 5.5 |

The MANOVA on source baselines (MPB, POM and terrestrial vegetation) isotopic signatures (Table 1) demonstrated significant differences between sources (F=11.21, df=4, p<0.001), sites (F=27.32, df=2, p<0.001), but no effect of vegetation type (F=2.03, df=4, p=0.094) and no interaction between site and vegetation type (F=0.05, df=2, p=0.943).

The MANOVA on potential prey isotopic signature demonstrated a significant effect of fish species (F=32.04, df=42, p<0.001), site (F=87.06, df=2, p<0.001) and vegetation type (F=5.10, df=4, p<0.001), but no interaction between site and vegetation type (F=0.234, df=2, p=0.790).

### 3.2. Fish diet partitioning

The diets of all three fish species were better explained by models on basal sources (MPB, POM and vegetation types) including both site and vegetation type (Table 2). Nevertheless, for *P. microps* the probability that a model only including site as a random factor could best explain the variations in diet remains high (weight = 0.413, Table 2).

**Table 2:** LOOic values of each mixing model tested with basal sources (microphytobenthos, particulate organic matter and terrestrial vegetation). High weight indicates high probability the model is the best explaining diet partition.

| Fish species | Model | LOOic | Std error LOOic | Weight |
| --- | --- | --- | --- | --- |
| *Dicentrarchus labrax* | Site +Vegetation |  |  |  |
|  | type | -125.1 | 25.8 | 1,0 |
|  | Site | -102.8 | 22.6 | 0,0 |
|  | Vegetation type | -7.6 | 24.7 | 0,0 |
|  | Null | 99.7 | 14.0 | 0,0 |
| *Pomatoschistus microps* | Site +Vegetation |  |  |  |
|  | type | 18.3 | 8.4 | 0.6 |
|  | Site | 19.0 | 8.3 | 0.4 |
|  | Vegetation type | 63.1 | 11.1 | 0.0 |
|  | Null | 70.2 | 9.0 | 0.0 |
| *Chelon ramada* | Site +Vegetation |  |  |  |
|  | type | 111.1 | 15.6 | 0.88 |
|  | Site | 115.1 | 14.2 | 0.1 |
|  | Vegetation type | 210.5 | 16.5 | 0.0 |
|  | Null | 221.9 | 16.3 | 0.0 |

Similar results were found for *D. labrax* and *P. microps* for the models based on potential prey (*Mysida, O. gammarellus, L. rugicauda* and *H. diversicolor*) (Table 3).

**Table 3:** LOOic values of each mixing model tested with potential prey as sources. High weight indicates high probability the model is the best explaining diet partition.

| Fish species | Model | LOOic | Std error LOOic | Weight |
| --- | --- | --- | --- | --- |
| *Dicentrarchus labrax* | Site +Vegetation type | -14.6 | 26.5 | 1,0 |
|  | Site | 22.5 | 22.7 | 0,0 |
|  | Vegetation type | 129.7 | 24.9 | 0,0 |
|  | Null | 239.6 | 14.9 | 0,00 |
| *P. microps* | Site +Vegetation type | 31.8 | 15.8 | 0.7 |
|  | Site | 33.3 | 14.3 | 0.3 |
|  | Vegetation type | 95.9 | 12.6 | 0.0 |
|  | Null | 100.9 | 12.0 | 0.0 |

The posterior estimates of overall diet proportions demonstrate that MPB and, in a lesser extent POM, are the main sources of organic matter contributing to *D. labrax* diet (Fig. 2). Site and vegetation type equally contribute to variation in *D. labrax* diet (site median standard deviation = 6.83 (95%CI = 1.65-17.84), vegetation type median standard deviation: 4.33 (95%CI = 1.09-16.17).

**Figure 2:**
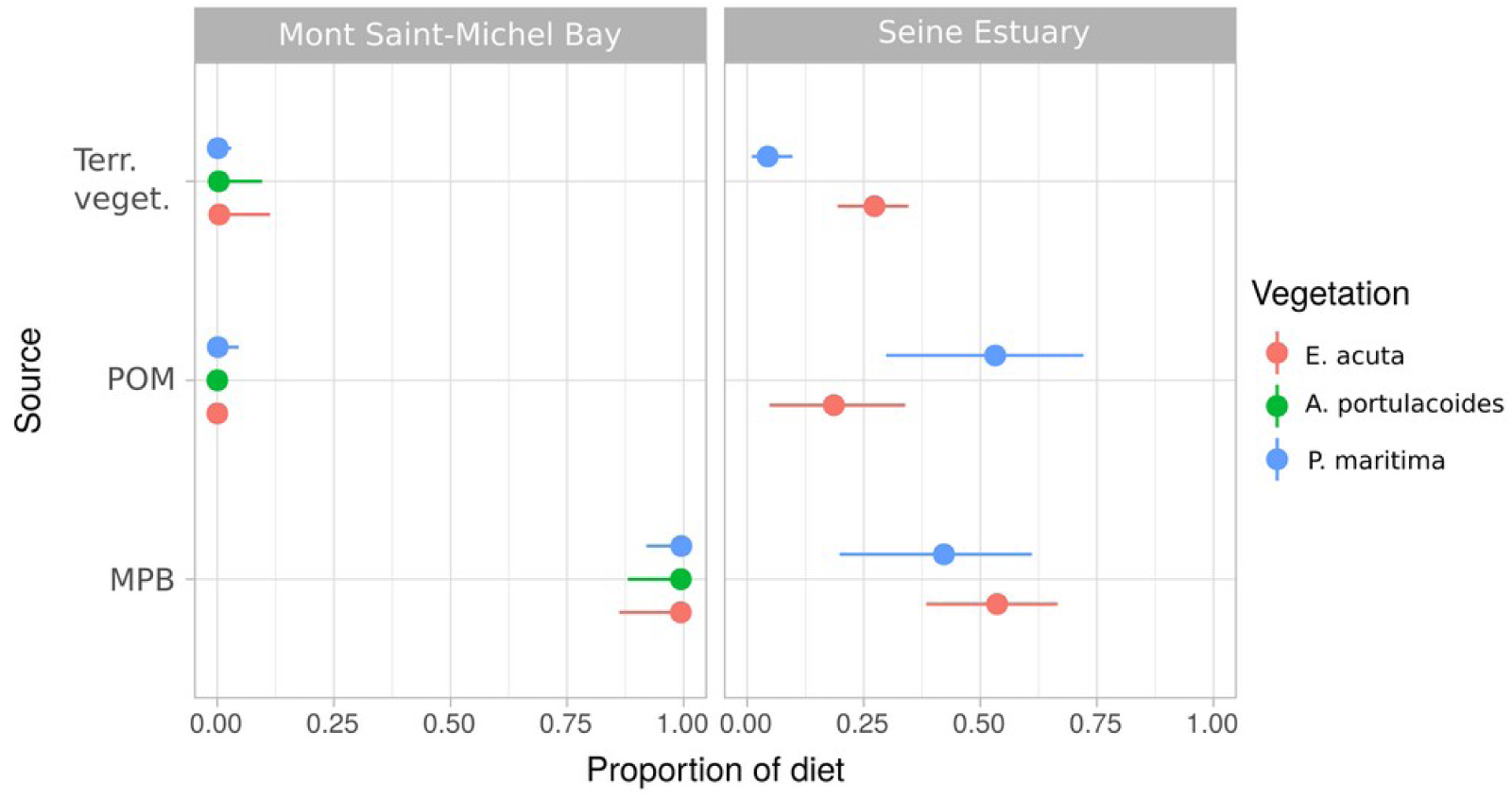
Posterior median (points) with 90% (thin lines) confidence intervals for the contribution of each source (MPB: microphytobenthos, POM: particulate organic matter, Terr. veg.: terrestrial vegetation) according to dominant vegetation in the sampling area for *Dicentrarchus labrax*.

For the sites located in the Mont-Saint-Michel Bay, the contribution of terrestrial sources to *D. labrax* diet is null whatever vegetation type the individuals were sampled in (Fig. 2 and Appendix B). In the Seine Estuary the contribution of terrestrial sources to *D. labrax* diet is 0.273 (95%CI = 0.194 – 0.361) for individuals collected in *E. acuta* areas and 0.044 (95%CI = 0.01 – 0.107) in *P. maritima* areas. Regarding aquatic sources, in the Mont-Saint-Michel Bay, the MPB is the main source for *D. labrax* (> 0.99) whatever vegetation type the individuals were sampled in (Fig. 2 and Appendix B). In the Seine estuary, POM and MPB both contribute to *D. labrax* diet in the same proportions (0.422-0.536, 95%CI =0.198-0.761) except in *E. acuta* areas where POM is lower (0.186, 95%CI = 0.048 – 0.365) (Fig. 2 and Appendix B).

The posterior estimates of overall diet proportions based on prey signatures show that *Mysida* was the main prey of *D. labrax* (contribution >0.5 in all cases, Fig. 3, Appendix C) but the contribution of *Orchestia gammarellus* increased up to 0.376 (95% CI: 0.276-0.475) in *E. acuta* dominated areas in the Seine Estuary (Fig. 3, Appendix C). Among the other aquatic prey, *Lekanesphaera rugicauda* was the second main source in areas dominated by *P. maritima* in both sites (Fig. 3, Appendix C), whereas it was *Hediste diversicolor* under a cover of *A. portulacoides* (Fig. 3, Appendix C). *O. gammarellus* contribution was significantly higher in *E. acuta* area in the Seine Estuary (Fig. 3, Appendix C).

**Figure 3:**
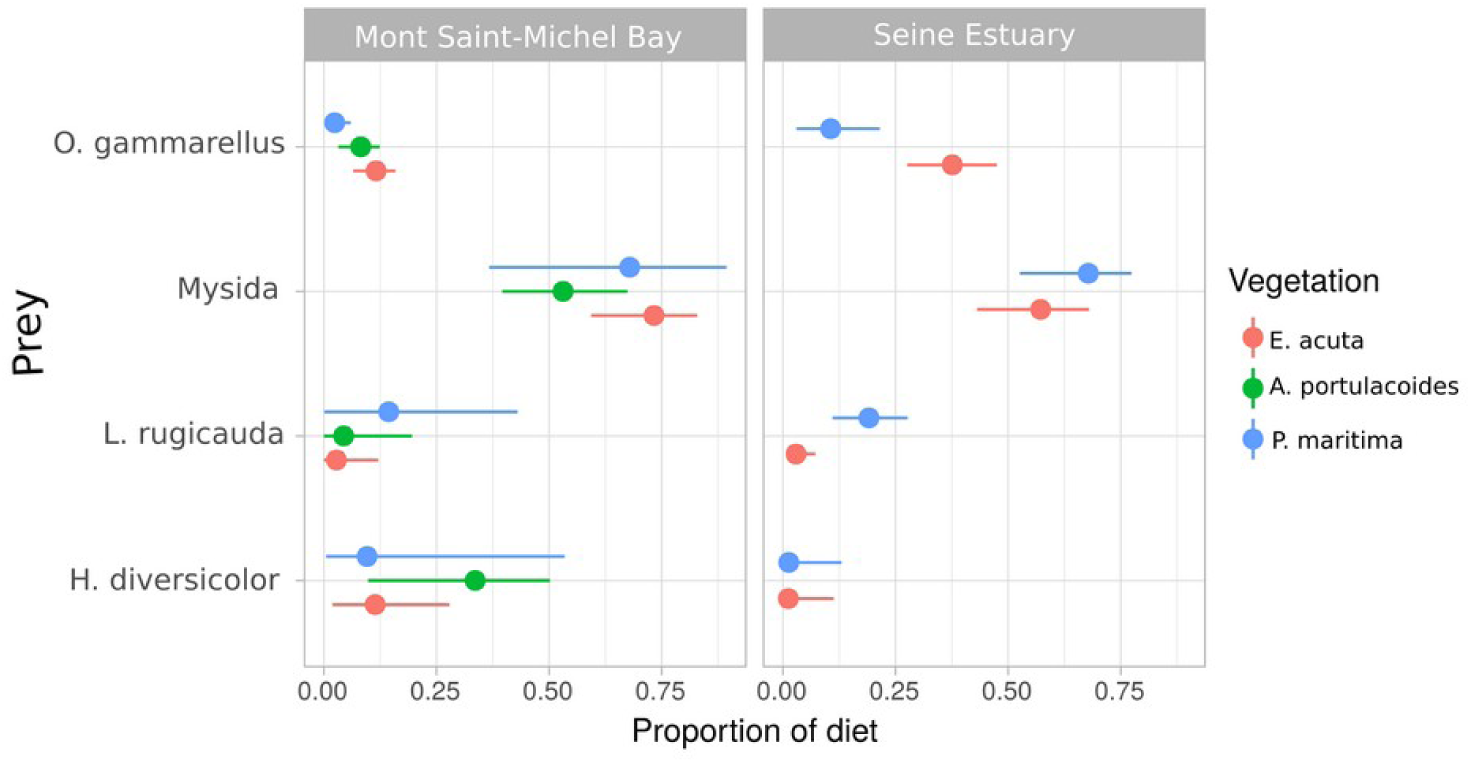
Posterior median (points) with 50% (thick segments) and 90% (thinner lines) confidence intervals for the contribution of each prey according to dominant vegetation in the sampling area for *D. labrax*. Site effect is included in the posterior estimate.

The posterior estimates of overall diet proportions show that *P. microps* main source of carbon was MPB in both sites, whatever vegetation type the fish were foraging in (Fig. 4, Appendix B). Site and vegetation type equally contributed to variation in *P. microps* diet (site mean standard deviation = 10.99 (95%CI = 1.07-19.23), vegetation type mean standard deviation: 6.19 (95%CI = 0.386-19.21). Nevertheless, no obvious difference in source contribution according to vegetation type was found (Fig. 4). For the areas of the Mont-Saint-Michel Bay, the contribution of terrestrial sources to *P. microps* diet was null for the three vegetation types (Appendix B)) while it was slightly superior in the Seine Estuary (<0.025, Appendix B).

**Figure 4:**
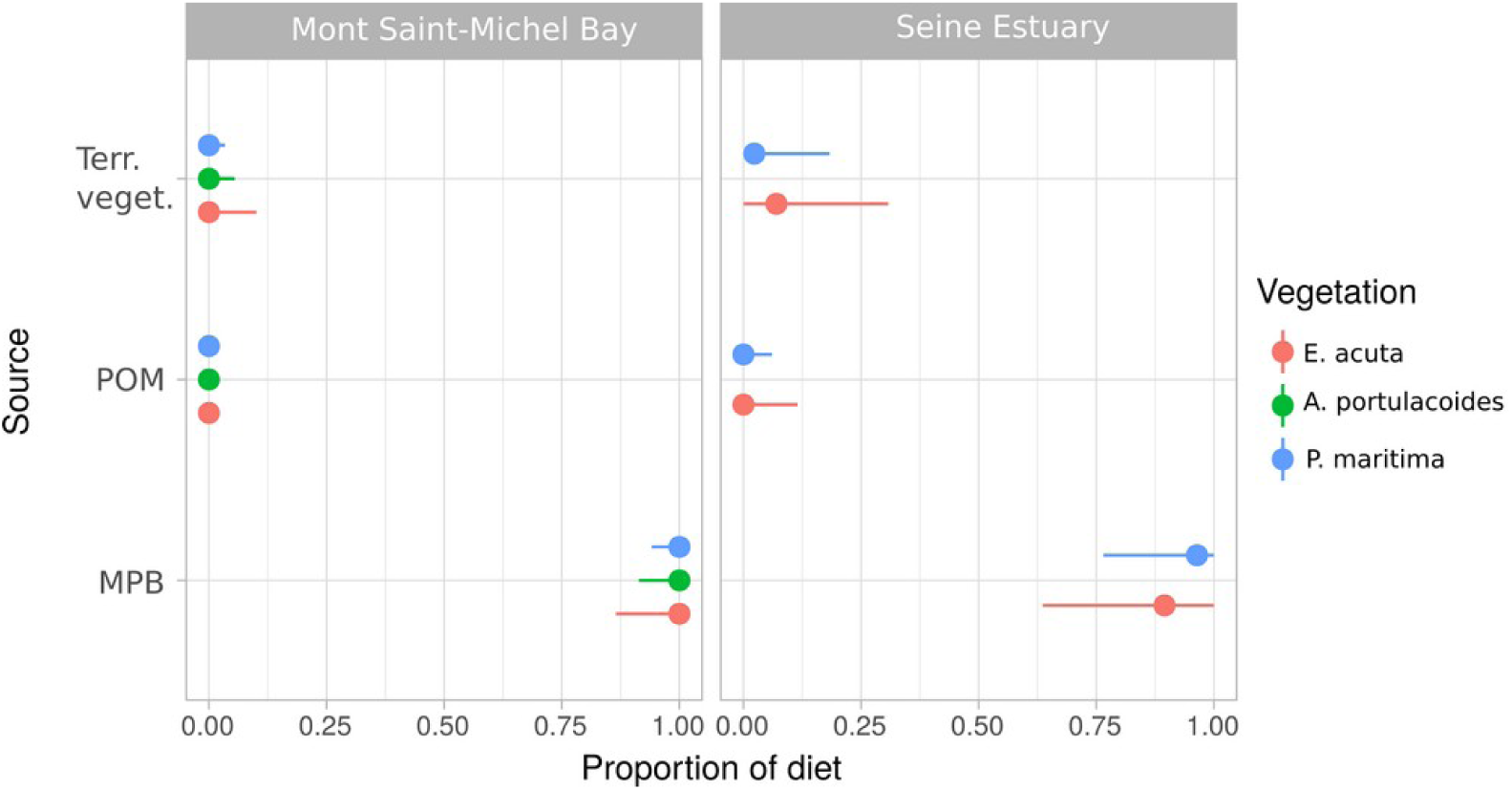
Posterior median (points) with 90% (thin lines) confidence intervals for contribution of each source (MPB: microphytobenthos, POM: particulate organic matter, Terr. veg.: terrestrial vegetation) according to dominant vegetation in the sampling area for *P. microps*.

The posterior estimates of overall diet proportions based on prey signatures show that *Mysida* was the main prey of *P. microps* (contribution >0.5 in all cases, Fig. 4, Appendix C) but the contribution of *H. diversicolor* was between 0.17 and 0.44 in the Mont-Saint-Michel Bay. No difference was found between the different vegetation types (Fig. 4, Appendix C).

The posterior estimates of overall diet proportions show that *C. ramada* main source of carbon were MPB in both sites and for all vegetation types (> 0.8, Fig. 5, Appendix B). In the Mont-Saint-Michel Bay terrestrial vegetation also slightly contributed to *C. ramada* diet (up to 0.17 for *A. portulacoides* Fig. 5, Appendix B). Site and vegetation type equally contributed to variation in *C. ramada* diet (site mean standard deviation = 8.62 (95%CI = 0.71-18.77), vegetation type mean standard deviation: 7.92 (95%CI = 1.03-18.22) (Fig. 4). Nevertheless, no difference was found between sites.

**Figure 5:**
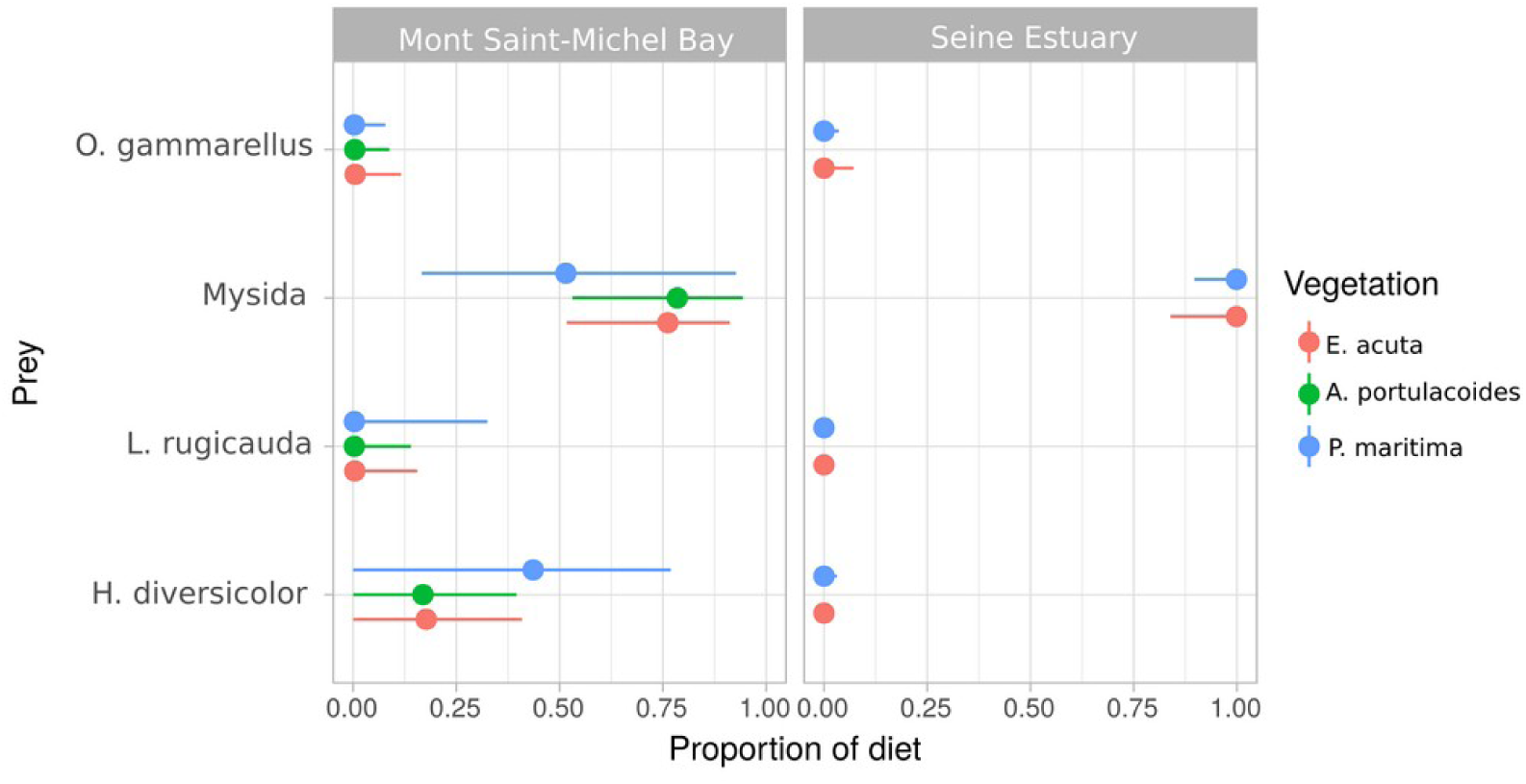
Posterior median (points) with 50% (thick segments) and 90% (thinner lines) confidence intervals for the contribution of each prey according to dominant vegetation in the sampling area for *P. microps*. Site effect is included in the posterior estimate.

**Figure 6:**
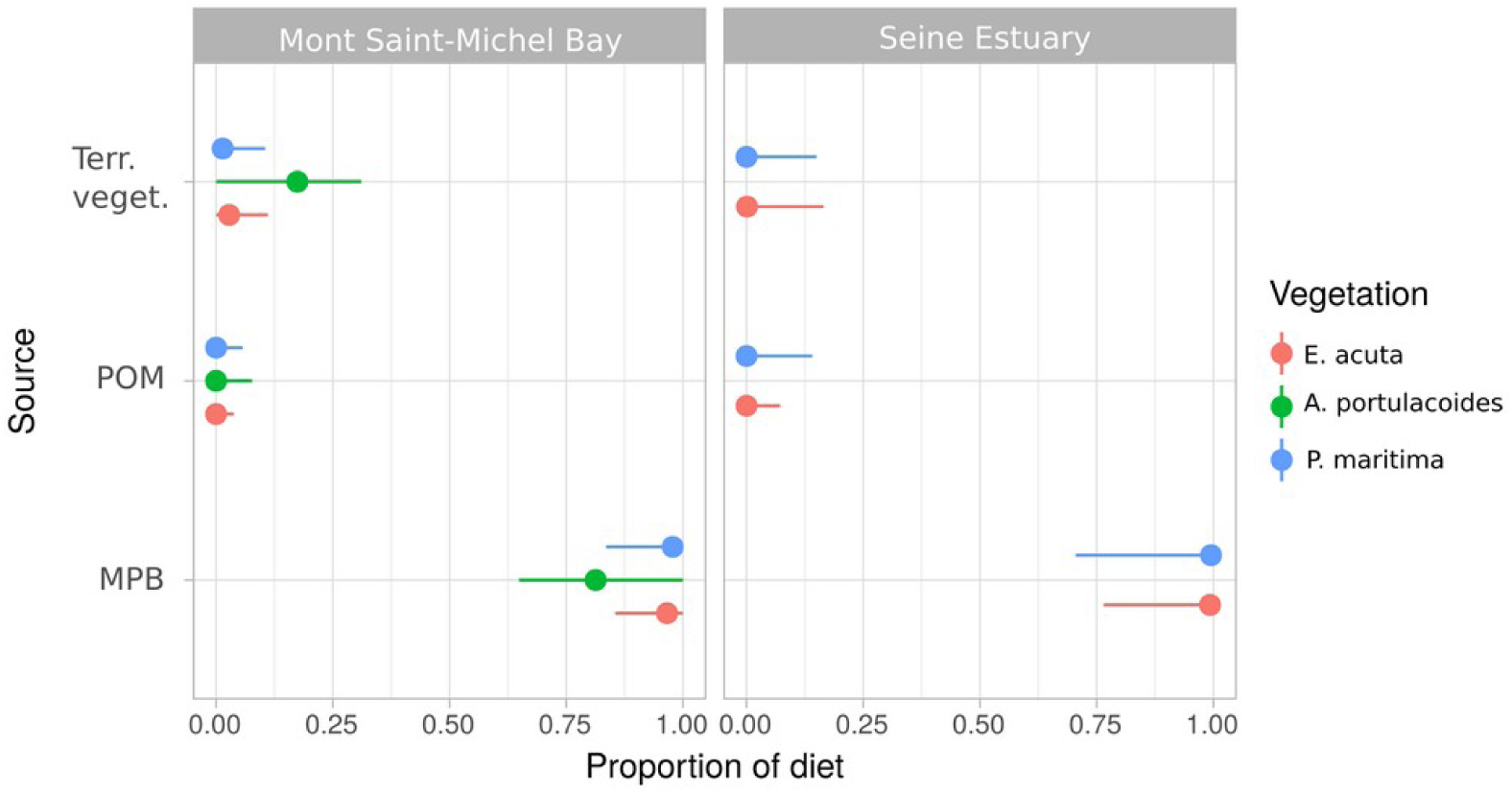
Posterior median (points) with 90% (thin lines) confidence intervals for contribution of each source (MPB: microphytobenthos, POM: particulate organic matter, Terr. veg.: terrestrial vegetation) according to dominant vegetation in the sampling area for *C. ramada*.

**Figure 7:**
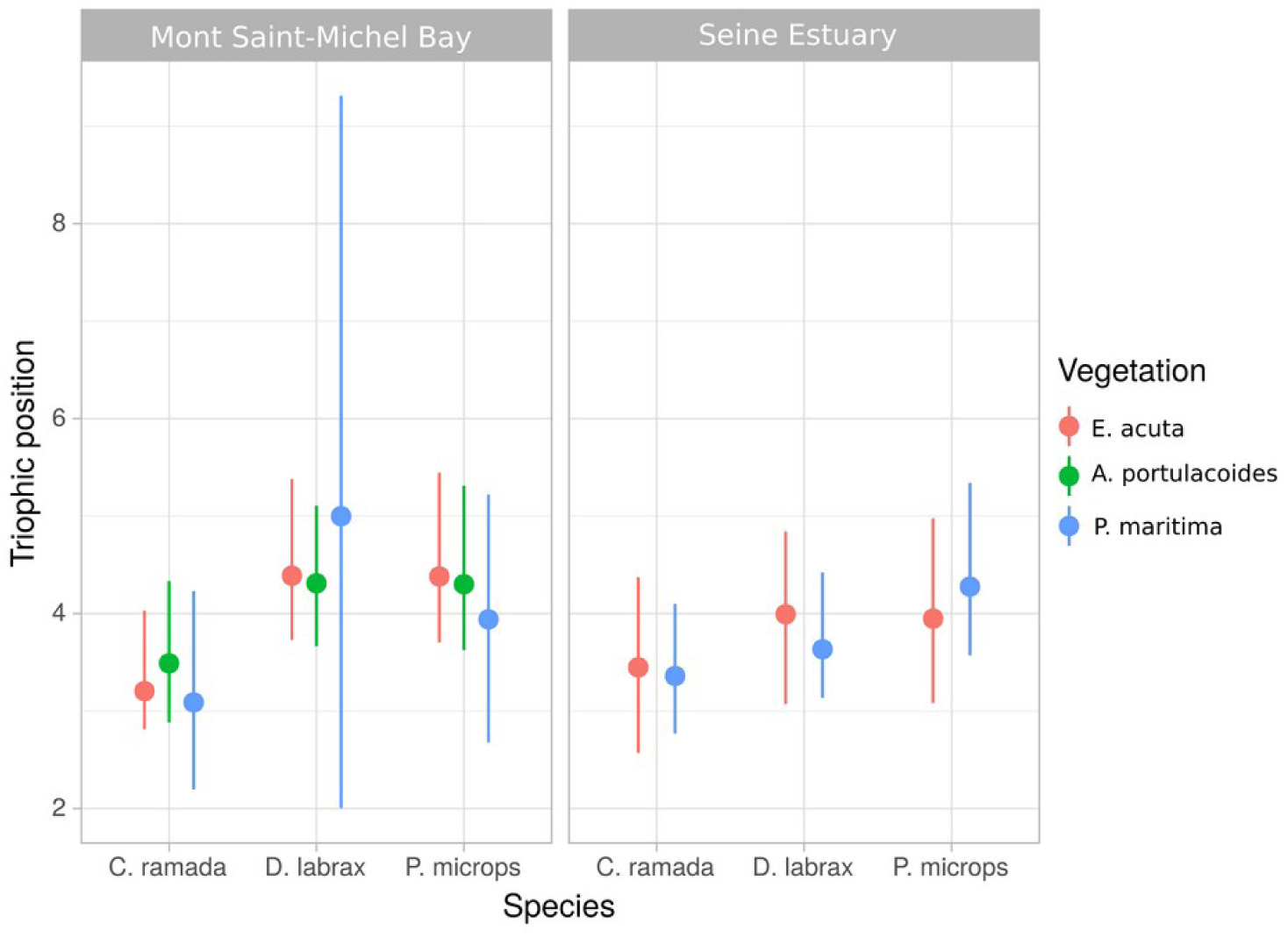
Median and 95%CI of posterior trophic position for each fish species according to vegetation type.

### 3.3. Isotopic niche

For the three fish species, the posterior means of the standard ellipse area were different between sites but not between vegetation types (Table 4). Values were consistently higher in the Seine Estuary than in the Mont-Saint-Michel Bay.

**Table 4:** Standard ellipse area and 95% confidence interval of each fish species per site and vegetation type. SE: Seine Estuary, MSMB: Mont-Saint-Michel Bay

| Fish | Vegetation type | Mont-Saint-Michel Bay |  | Seine Estuary |  |
| --- | --- | --- | --- | --- | --- |
|  |  | Standard Ellipse Area (‰<sup>2</sup>) | 95%CI | Standard Ellipse Area (‰<sup>2</sup>) | 95%CI |
| *Dicentrarchus labrax* | *Elytrigia acuta* | 0.508 | 0.283 – 0.879 | 1.110 | 0.743 – 1.854 |
|  | *Atriplex portulacoides* | 0.383 | 0.248 – 0.601 | 1.132 | 0.736 – 1.657 |
| *Pomatoschistus microps* | *Elytrigia acuta* | 0.408 | 0.168 – 1.061 | 1.378 | 0.600 – 3.848 |
|  | *Atriplex portulacoides* | 0.408 | 0.165 – 1.036 | 1.678 | 0.681 – 4.546 |
| *Chelon ramada* | *Elytrigia acuta* | 3.084 | 1.694 – 5.326 | 4.706 | 4.603 – 7.235 |
|  | *Atriplex portulacoides* | 1.788 | 0.969 – 3.185 | - | - |
|  | *Puccinellia maritima* | 6.318 | 2.008 – 16.123 | 5.761 | 2.912 – 11.539 |

### 3.4. Fish trophic position

Posterior estimate of trophic position of each fish species was not different between vegetation types (probabilities of significant differences consistently inferior to 0.5, Appendix B, Fig. 5). On the other hand, systematic high probabilities of difference were observed when comparing sites and fish species trophic position between each other. *C. ramada* had a high probability of having lower trophic position than *D. labrax* and *P. microps* and, for the same fish species, individuals from the Seine Estuary had higher trophic positions than individuals from the Mont-Saint-Michel Bay (Appendix B, Fig. 5).

## 4. Discussion

To the best of our knowledge, this study is the first to investigate the effect of vegetation type (resulting from invasion by *E. acuta* and sheep grazing) on salt-marsh nursery function for juvenile fish using stable isotopes. We found a clear difference in the contribution of main carbon sources to *D. labrax, P. microps* and *C. ramada* diets. Nevertheless, vegetation type had a weak influence to these contributions except for *D. labrax*, in the Seine estuary only. The most striking and unexpected result of our study was actually the loss of sea bass nursery function formerly provided by salt marshes in the Mont Saint-Michel Bay (MSMB). Interestingly, we could not find any difference of trophic position or isotopic niche between the different vegetation types for any of the three fish species studied.

Results of Bayesian mixing models suggest that the contribution of carbon sources to juvenile sea bass did not vary along with salt-marsh vegetation where fish are foraging (i.e. >99% microphytobenthos (MPB)) in the MSMB. MPB still dominated in the Seine estuary but other sources (POM and terrestrial vegetation) significantly contributed to the sea bass diet, in proportions differing according to the surrounding plant species. The mixing model based on preys revealed that *Mysida*, a planktonic taxa, appears to be the main prey of juvenile sea bass in both sites, but secondly completed by *O. gammarellus* and *L. rugicauda* in the Seine estuary and *H. diversicolor* in the MSMB, thereby confirming the different influences of C sources in the two sites.

From these results, one can state that in the Seine estuary, the salt-marsh C resources have a significant contribution to the diet of *D. labrax* juveniles reaching almost 30% in areas dominated by *E. acuta*. On the opposite, the terrestrial contribution of salt marshes to *D. labrax* diet was almost null in the MSMB. In the MSMB, Laffaille et al. (2001) also identified *Mysida* as the main prey of sea bass juveniles entering salt marshes but *O. gammarellus* was the main prey in the stomach of individuals leaving salt marshes during ebb. In 2005, Laffaille et al. demonstrated, based on stomach content analyses, that in the MSMB sea bass juveniles were consuming less *O. gammarellus* in areas invaded by *E. acuta*. Laffaille et al. (2005) estimated the numerical contribution of *O. gammarellus* to the diet of sea bass juveniles as 58 % (73% in biomass) in uninvaded areas and 28 % (40% in biomass) in invaded areas. They hypothesized that such change in diet was due to a decrease in accessibility of terrestrial preys (here *O. gammarellus*) because of changes in vegetation structure (increased stem density was seen a physical obstruction for fish to reach amphipods). Global availability was also reduced, but considered by the authors still high enough to not act a limiting factor (see also Mantzouki et al., 2012). Our results suggest that, about 10 years after Laffaille et al. study (2005), the contribution of terrestrial sources to *D. labrax* diet decreased down close to zero. We hypothesise that the access to prey was modified not only by changes in vegetation, but also by changes in the morphology of creeks, resulting from increasing nitrogen inputs (also probably the main factor explaining the spread of *E. acuta* in the Saint-Michel Bay: Valéry et al. (2017). In a recent study, Nelson et al. (2019) indeed demonstrated that the nutrient enrichment of waters entering salt marshes triggers a cascade of effects and positive feedback resulting in geomorphological changes (creek deepening and enlargement with cliff-like bank: see Deegan et al., 2012 for a detailed description of processes) decoupling the high-marsh ecosystem from the creek ecosystem. Similarly to our results on sea bass in the MSMB, the authors observed a lower contribution of terrestrial prey to mummichog (*Fundulus heteroclitus*) diet. This hypothesis is reinforced by the situation in the Seine estuary where *E. acuta* is not invasive and where sea bass juveniles appear more dependant to the saltmarsh resources (*O. gammarellus* contributed up to 38% in diet partitioning). In this site, *D. labrax* diet is varying with vegetation type. We are aware that this interpretation implies that site fidelity exists for juveniles of sea bass. Despite several studies demonstrated that sea bass are site (and even creek) dependent (Green et al., 2012; Joyeux et al., 2017; Vinagre et al., 2011), the question is still under debate and needs to be investigated in our study sites.

The differences in terrestrial C contributions between the two sites can also be the result of different frequency of access to salt marshes for fish. The two sites indeed strongly differ in tidal amplitude (7m in the MSMB vs 15m in the Seine estuary) and high water slack duration (null in the MSMB vs. around two hours in the Seine estuary because of frequent embankments). As a consequence, the percentage of tides during which fish are able to enter into salt-marsh creeks are estimated to 43% over one year in the MSMB (Laffaille et al., 2001) and to 80% in the Seine estuary (Duhamel comm.pers.). One can thus hypothesize that such a difference in possibilities for fish to exploit salt-marsh creeks for feeding would have a significant effect on their isotopic signature.

*Pomatoschistus microps* presented a similar pattern of diet shift. MPB contributed to 100% of its diet in the MSMB, whatever the surrounding vegetation type was, and >90% in the Seine estuary where the terrestrial vegetation contributed to no more than 2%. Dominat preys were *Mysida* in both sites (100% in Seine estuary) and secondly an Annelid (*H. diversicolor*) in the MSMB (reaching 44% in *P. maritima* areas). These preys are commonly described for this species in other European sites in where it is described a opportunistic benthic-feeding carnivore (Magnhagen, 1986; Magnhagen and Wiederholm, 1982; Salgado et al., 2004). Based on stomach contents, dominant preys are amphipods (*C. volutator*) in Magnhagen and Winderholm 1982, Magnhagen 1986 but also annelids in Gee et al. (1985) and Salgado et al. (2004). Interestingly, Laffaille et al. (2005) revealed that, in the MSMB, another species of *Pomatoschistus* (*P. minutus*) switched its diet in invaded areas (by *E. acuta*) from *O. gammarellus* to *H. diversicolor* (the numerical contribution of *O. gammarellus* was 72% - 76% in biomass-in uninvaded areas and 10 % - 15% in biomass - in invaded areas when the numerical contribution of *H. diversicolor* was 3% - 13% in biomass – in uninvaded areas and 18% - 69% in biomass - in invaded areas). As *P. microps* is considered a semi-resident species, it was expected to be the fish most influenced by changes in salt-marsh vegetation. But in the MSMB, like for sea bass, the function of feeding (*P. microps* was represented by both adults and juveniles in this study) could have been impacted by the dominance of invasive *E. acuta*. In the Seine estuary, the dominance of *Mysida* in the goby diet is more complex to interpret. The very low contribution of terrestrial vegetation confirms this, and tends to demonstrate that salt marshes were less important for this species, as compared to the sea bass for instance. The comparative approach used here consequently reveals that resident species of a single taxa (here fish) may react very differently to important changes in their usual habitat, or larger surfaces used for foraging (here surrounding vegetation).

As expected, juveniles of the mullet *C. ramada* derived most of their carbon from benthic sources, and particularly MPB. Similar results were reported by Lebreton et al. (2011) using stable isotopes. These authors confirmed that juvenile mullets exploiting salt marshes are limno-benthophagous. Nevertheless, the uncertainty regarding the contribution of each source was very high in our study. This result suggests that this species uses several salt marshes with different isotopic signatures to forage. This hypothesis is corroborated by the fact that *C. ramada* had the widest isotopic niche despite its lower trophic position (*C. ramada* usually feeds upon MPB and meiofauna consumers of MPB, see Carpentier et al., 2014 and Lebreton et al. 2011) compared to *D. labrax* and *P. microps*.

The trophic position of the three studied fish species was not different according to the vegetation type or site. This result is not surprising as we found a very weak contribution of terrestrial sources to fish diet in the MSMB. In the Seine estuary, despite the higher contribution of terrestrial sources to fish diet, the trophic position of fish did not vary with vegetation type.

No significant difference in isotopic niche was found according to vegetation type for *D. labrax*. This demonstrates that the changes induced by the invasive plant *E. acuta* on the aquatic food web remained weak. The isotopic niches of *P. microps* and *C. ramada* were not different according to vegetation types. As for trophic position, these results are in accordance with the diet partitioning results. Indeed, given the weak contribution of terrestrial sources to *P. microps* and *C. ramada* diet, variations in their niche breadth along changes in surrounding vegetation were very unlikely.

It is finally interesting to note that we found a strong, unexpected, site effect on all characteristics of the food web. All best diet partitioning models indeed included site as a predictive variable, with site and vegetation type equally contributing to variation in fish diet. Nevertheless, the large uncertainty on the contribution of the two variables renders any comparison hard. The contribution of terrestrial sources to predatory fish diet was much higher in the Seine Estuary. We also found the isotopic niche breadth of all three species significantly higher in the Seine Estuary. At a large spatial scale, a recent review by Ziegler et al. (2019) emphasised on the key role of tidal amplitude in structuring food webs. We state that a similar process (i.e. tides regulate the access to salt marshes for transient estuarine fishes) can be observed within the same biogeographic area. Particularly we hypothesise that the importance of nursery function of macrotidal salt marshes in Western Europe would be more diverse than expected given the large inter-site variability and overall low terrestrial contribution to fish diet we found in this study. Future large-scale studies assessing actual salt-marsh contribution to aquatic food webs, for instance along a gradient of tidal amplitude, are needed to test this assumption.

## Acknowledgements

We would like to thank the Programme Seine aval which financed this study included in the DEFHFIS program. We also thank the Agence de l’eau Loire-Bretagne (PEPPS project) and the Région Bretagne (SAD) for supporting D. Lafage. We also want to thank the Plateforme de Spectrométrie isotopique of the LIENSs laboratory of La Rochelle for the isotopic analyses.

## Appendix A

Mean and standard deviation of δ13C and δ15N of prey and sources per site and vegetation type

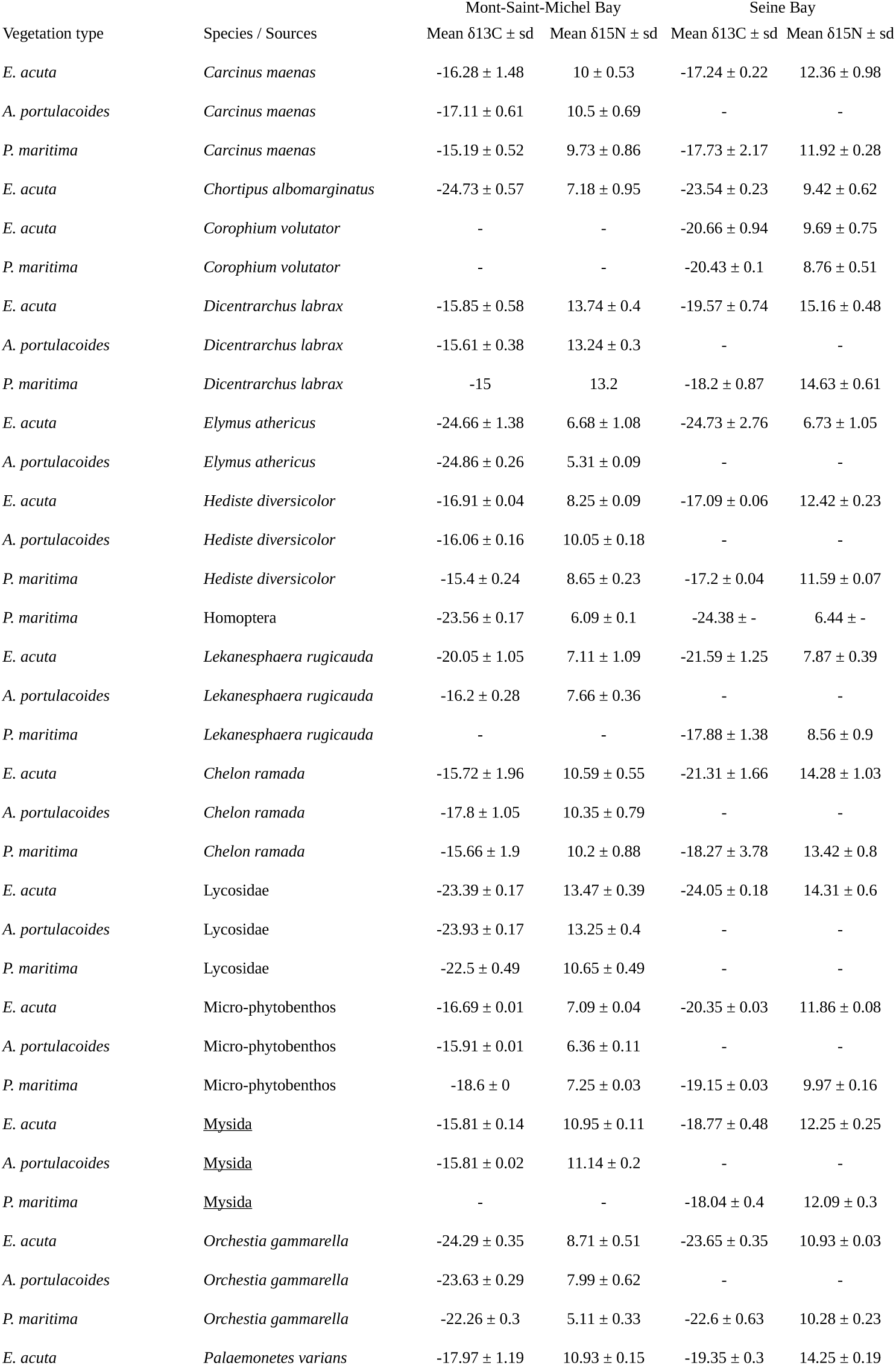

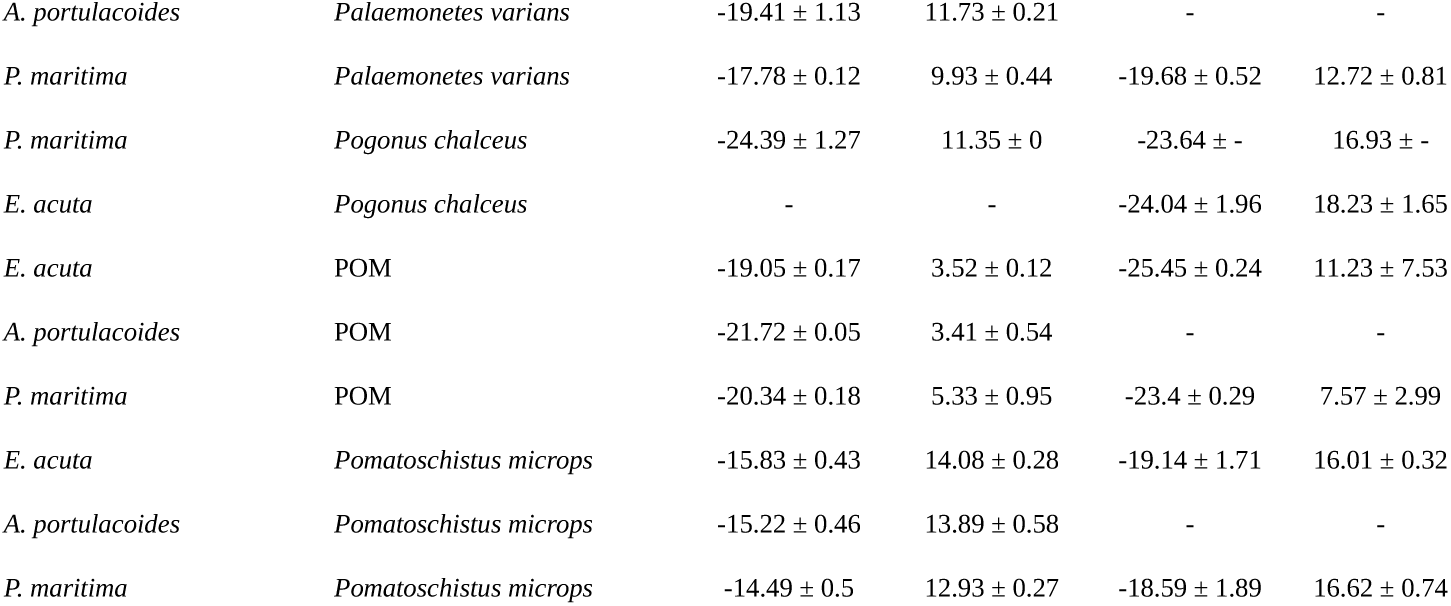

## Appendix B

Results of MixSIAR models for diet partitionning of *Dicentrarchus labrax* (*D. labrax*), *Pomatoschistus microps* (*P. microps*) and *Chelon ramada* (*C. ramada*) based on 3 sources (MPB: microphytobenthos, POM: particulate organic matter, Terr. veg.: terrestrial vegetation). Posterior median are given with 95% confidence intervals.

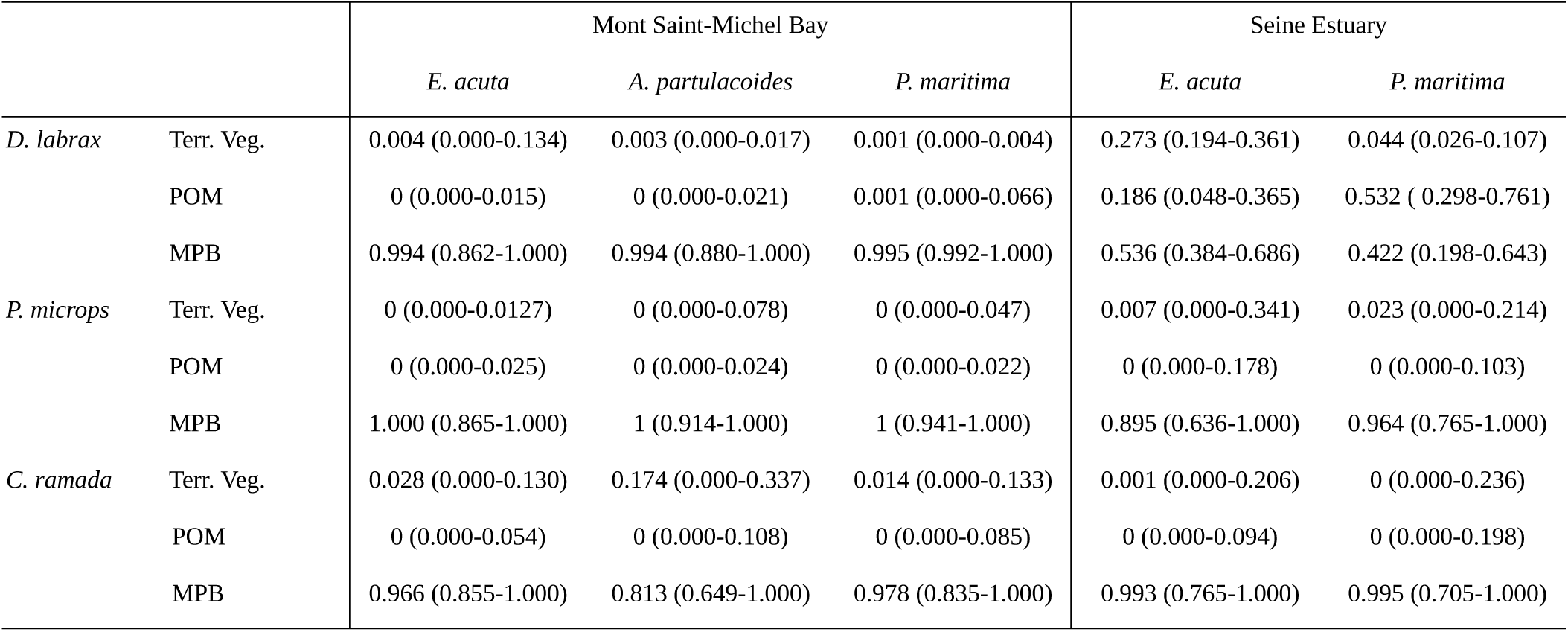

## Appendix C

Results of MixSIAR models for diet partitionning of *Dicentrarchus labrax* (*D. labrax*) and *Pomatoschistus microps* (*P. microps*) based on 4 prey MPB. Posterior median are given with 95% confidence intervals.

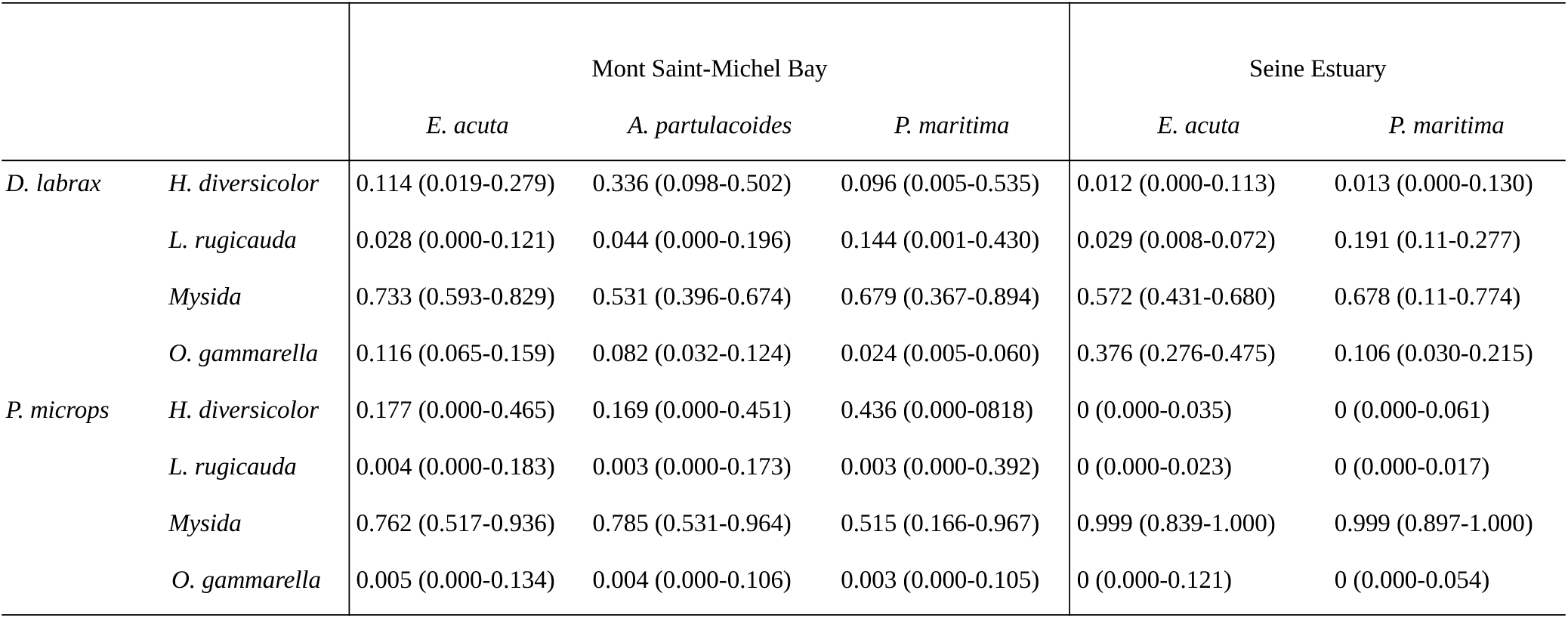

## Appendix C

Probability of trophic position of a fish foraging in a given vegetation type and site to be inferior or equal to another one. SE: Seine Estuary, MSMB: Mont-Saint-Michel Bay.

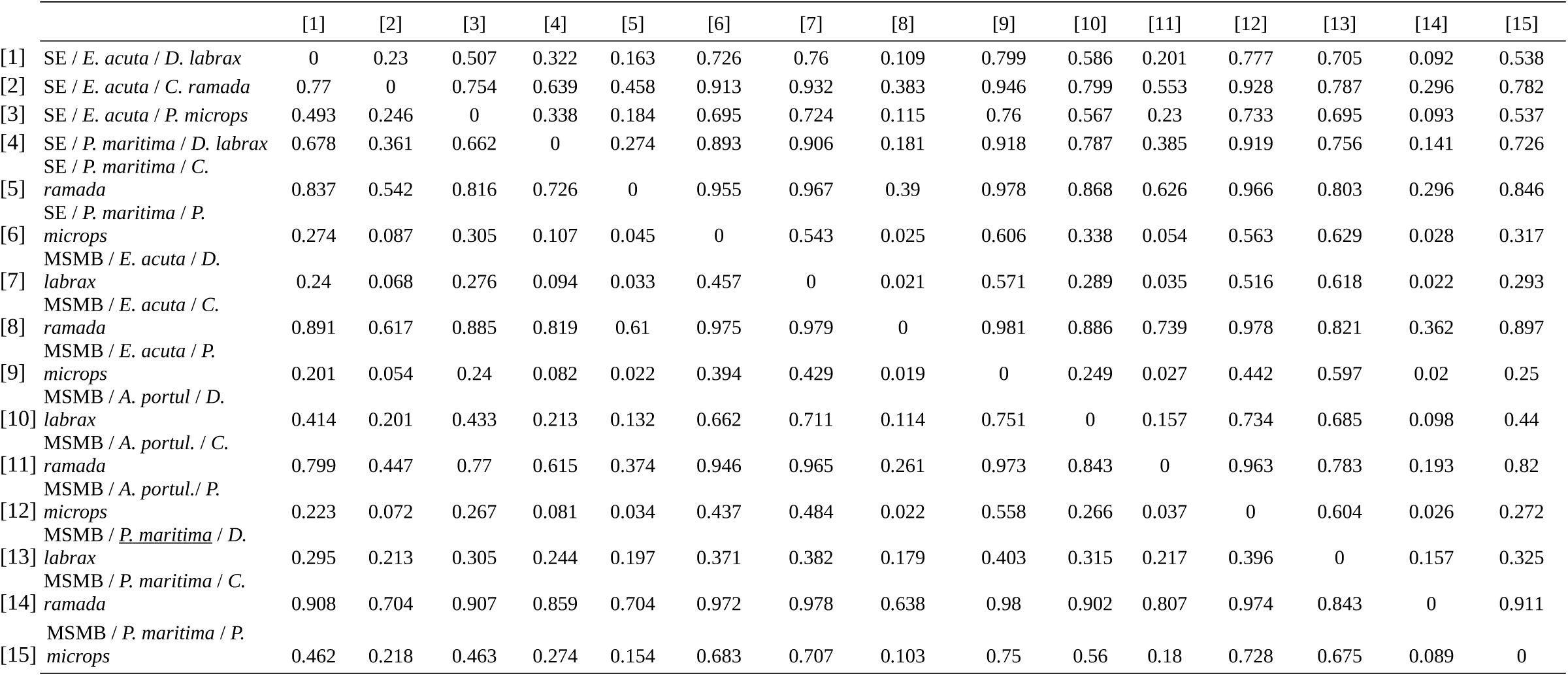

